# Flexible integration of natural stimuli by auditory cortical neurons

**DOI:** 10.1101/2024.04.18.590153

**Authors:** Grace Wan Yu Ang, Claudia Clopath, Andriy S. Kozlov

**Affiliations:** Department of Bioengineering, Imperial College London

## Abstract

Neurons have rich input-output functions for processing and combining their inputs. Although many experiments characterize these functions by directly activating synaptic inputs on dendrites *in vitro*, the integration of spatiotemporal inputs representing real-world stimuli is less well studied. Using ethologically relevant stimuli, we study neuronal integration in relation to Boolean AND and OR operations thought to be important for pattern recognition. We recorded single-unit responses in the mouse auditory cortex to pairs of ultrasonic mouse vocalization (USV) syllables. We observed a range of integration responses, spanning the sublinear to supralinear regimes, with many responses resembling the MAX-like function, an instantiation of the OR operation. Integration was more MAX-like for strongly activating features, and more AND-like for spectrally distinct inputs. Importantly, single neurons could implement more than one integration function, in contrast to artificial networks which typically fix activation functions across all units and inputs. To understand the mechanism underlying the flexibility and heterogeneity in neuronal integration, we modelled how dendritic properties could influence the integration of inputs with complex spectrotemporal structure. Our results link nonlinear integration in dendrites to single-neuron computations for pattern recognition.

**Significance statement:** Sensory neurons compute over their inputs, combining them in ways to achieve selectivity and invariance for pattern recognition. Using real-world stimuli, we show that single cortical neurons are flexible, being capable of implementing more than one computation, unlike artificial neural-network units with fixed activation functions. We investigate this flexibility by modeling how synaptic activation patterns of real-world stimuli affect dendritic integration and the resultant neuronal computation. Our work bridges the gap between biophysical mechanisms and computation, linking neuronal input integration to pattern recognition.

## 1 Introduction

Network computations are shaped by how single neurons integrate their inputs to produce output signals. The input-output functions of neurons have been shown to be highly diverse [Gidon et al., 2020, Lafourcade et al., 2022, Tran-Van-Minh et al., 2016] and dynamic [Młynarski and Hermundstad, 2021], enabling the network to perform rich and complex tasks [Wybo et al., 2023, Geadah et al., 2023]. To investigate neuronal integration, numerous studies have stimulated dendrites with well-controlled patterns of synaptic inputs *in vitro* [Schiller et al., 1997, 2000, Branco and Häusser, 2011, Tran-Van-Minh et al., 2016, Gidon et al., 2020] or measured responses to simple stimuli [Bölinger and Gollisch, 2012, Deny et al., 2017]. While these approaches have provided essential insights into the rules and biophysical mechanisms governing the integration of inputs, there is a gap in understanding how neurons combine natural, complex stimuli *in vivo*.

Single-neuron transfer functions can be approximated as logical operations used in computations [Kouh and Poggio, 2008, Riesenhuber and Poggio, 1999]. Among these, the OR and AND logical operations are thought to be particularly relevant for pattern recognition in sensory processing. The AND operation increases feature complexity for selectivity by summing over inputs encoding simpler features. On the other hand, MAX-pooling, an instantiation of the logical OR operation, confers invariance as the neuron’s response is primarily determined by its strongest input. These feature recombination functions were first demonstrated in the cat visual cortex [Lampl et al., 2004] and more recently in the central auditory system of songbirds [Kozlov and Gentner, 2014], with the latter study employing ethologically relevant stimuli to probe pooling responses.

In this study, our goal is to bridge the two complementary perspectives on neuronal input integration: the bottom-up approach focusing on the biophysical mechanisms that transform synaptic inputs to outputs, and the top-down perspective on computations performed by neurons to achieve selectivity and invariance. By recording extracellular single unit spikes in mouse auditory cortex, we measure integration responses to pairs of ultrasonic mouse vocalization (USV) syllables. We then model the integration of these ethologically relevant stimuli using a simple neuron model. This allows us to explore how the spatiotemporal pattern of synaptic inputs arriving onto dendrites shape the neuron’s resultant feature integration operation.

## 2 Materials and Methods

### 2.1 Surgical preparation

All procedures were carried out under the terms and conditions of licences issued by the UK Home Office under the Animals (Scientific Procedures) Act 1986. Extracellular recordings were made in the auditory cortex of adult female C57BL/6 mice (N=21, aged 6 to 11 weeks). Animals were anaesthetized using a mixture of fentanyl, midazolam and medetomidine (0.05, 5 and 0.5 mg/kg, respectively). A small craniotomy was made over the auditory cortex, at the caudal end of the left squamosal suture, centered 1.5 mm rostral to the lamdoid suture. The dura was removed and the surface of the brain was coated in a layer of silicon oil. The head was fixed to a rigid clamp and the mouse was transferred to an acoustic chamber.

### 2.2 Electrophysiological recording

Extracellular signals were recorded using a single-shank 32-channel silicon multi-electrode probe (Neuronexus), and acquired at a sample rate of 30 kHz (Neuronexus Smartbox Pro). The shank was lowered slowly and allowed to settle for at least 30 min before each recording. Spikes were extracted using the automatic spike sorting algorithm Kilosort3 (https://github.com/MouseLand/Kilosort) and manually curated with Phy (https://github.com/cortex-lab/phy). Units were considered well-isolated single units if they showed clear refractory periods in their autocorrelogram and had fewer than 1% spikes within a 1-ms interspike interval. Only stable units that exhibited little mechanical drift and that showed robust responses to repetitions of mouse USVs were identified for subsequent stages of the experiment.

### 2.3 Stimuli

Stimuli were delivered through an Avisoft UltraSoundGate Player (Avisoft Bioacoustics) connected to a PC. The sound level was adjusted such that the peak intensty did not exceed 85 dB SPL. To identify syllables for the summation index (SmI) measurements, a USV stimulus consisting of concatenated mouse vocalization song bouts [Lu et al., 2023] was presented for 10-15 repetitions. Syllables, approximately 100 ms in duration (based on the average duration of individual sylables) eliciting peaks in the PSTH were clipped and a 5-ms cosine ramp was applied to the start and end of each clip. When superimposing syllables to measure the combined stimulus response, syllables were combined with different temporal alignments, shifting from -20 ms to + 20 ms in 10 ms increments. This was to ensure that the MAX response was not an artifact from stimuli that were misaligned in time, which would produce a smaller than expected response. For cross-adaptation stimuli, the deviant syllable was concatenated with the train of adapting syllables with a 15-ms silence between each syllable. All stimuli were presented for 10-15 repetitions and in a randomized order, with a 1-s interval between each presentation. Routines to create and manipulate the stimuli were written in Python; pitch-shifting was performed using the Librosa library.

### 2.4 Analyses

#### Summation Index (SmI)

The summation index described in previous studies was used as a metric to quantify feature recombination functions [Sato, 1989, Lampl et al., 2004, Kozlov and Gentner, 2014]:

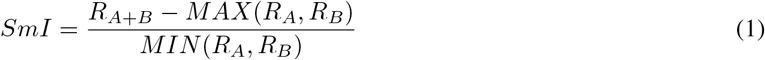

where *R*_*A*+*B*_ is the response to the combined stimulus (superimposed syllables) and *R*_*A*_ and *R*_*B*_ the responses to individual stimuli. Stimuli pairs for SmI calculations were included in the analyses if the response to each syllable was significantly greater than spontaneous activity (500 ms before stimulus onset). When individual responses did not meet this criterion, the stimulus pair was included if the firing rate to the combined stimulus was at least 25% higher than the dominant response to individual stimuli.

#### Pitch selectivity

When responses to *n* pitch-shifted versions of a syllable were recorded, pitch selectivity, *S*, was calculated using a metric previously described in Steadman and Sumner [2018]:

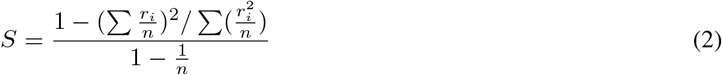

where *r*_*i*_ is the firing rate to the *i*^*th*^ pitch-shifted version of the syllable.

#### Syllable dissimilarity

For each pair of syllables used to measure the neuron’s SmI, we computed two distance metrics: (i) spectrogram distance and (ii) spectral feature difference. The spectrogram distance was the euclidean distance between flattened spectrograms of syllables. Spectrograms were generated using the stft function in Librosa, with *nfft* = 1024 and a Hanning window with 75% overlap. The second metric, the spectral feature difference, was the euclidean distance between feature vectors consisting of spectral mean, quartiles, and standard deviation of the sound waveform. These spectral characteristics were calculated using the BioSound class from the soundsig package.

To calculate the temporal correlation between syllables, we computed the Pearson correlation coefficient between their respective power profiles. These profiles provide a temporal representation of the overall energy of the syllable at each time point, and were obtained by averaging the spectrogram across frequency bins for each sound signal. Temporal correlation and spectral features were computed for the syllable segment that was within a 55-ms window of the peak response (averaged over repetitions) to the combined stimulus.

#### Regressing SmI on response and feature properties

A mixed-effects linear regression was fit to account for the contribution of individual factors to SmI. For syllable pair *i* presented to neuron *j*, this was described as:

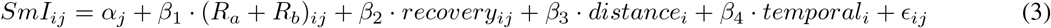

where the fixed effects, each associated with a coefficient *β*, are the predictors in the model: (i) sum of responses to individual syllables, (*R*_*a*_ + *R*_*b*_)_*ij*_, (ii) recovery from cross-adaptation *recovery*_*ij*_, (iii) spectral feature distance *distance*_*i*_ and (iv) syllable temporal correlation *temporal*_*i*_. The residual errors *ϵ*_*ij*_ are assumed to follow a normal distribution: *ϵ*_*ij*_ *∼ 𝒩* (0, *σ*^2^). The varying intercept *α*_*j*_ accounts for neuron-specific (random) effects and is assumed to be drawn from a distribution: 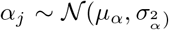, where *μ*_*α*_ and 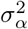 are the mean and variance of the random effects, estimated from the data. SmI values were converted to a normal distributions by a Box-Cox transformation (exponent *λ* = 0.38). The model was fit using the lmerTest function in R, and p-values for predictor coefficients were calculated using Satterthwaite’s approximation.

Marginal *R*^2^ values quantifying the proportion of variance explained by the predictors were calculated using the `partialR2` function in R for mixed-effects models. All statistical analyses were performed in R.

### 2.5 Neuron model

We simulated the integration of complex USV stimuli with a simple neuron model connected to 10 cylindrical dendritic subunits. The sound stimulus was first converted into spike trains which were inputs to the dendritic tree of a central auditory neuron. Code for the model is available at https://gitlab.com/kozlovlabcode/flex_integration.git.

#### 2.5.1 Stimulus encoding

To encode the sound stimulus into spikes, we used a neuron model based on methods described in Goodman and Brette [2010] for simulating cochlear filtering. Sounds were passed through a bank of 3000 gammatone filters evenly spaced on the Equivalent Rectangular Bandwidth (ERB) scale. The center frequencies of these filters ranged between 30 - 120 kHz, which encompassed the frequency content of USV syllables used in the experiments. The filtered sounds, *x* (sound pressure in pascals), were subjected to half-wave rectification and compression to generate a current 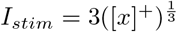. This current, together with noise 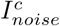, was fed as an input 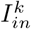 to a point neuron. The evolution of the membrane potential,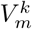, where the superscript *k* refers to either the stimulus-encoding neuron (denoted as *c*) or a neuronal compartment (as detailed in the following section), follows the dynamics of a leaky integrate-and-fire model with the standard form:

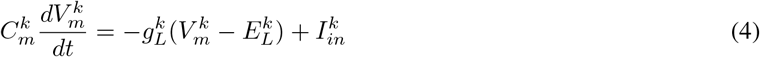

where 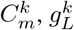 and 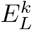 are the membrane capacitance, leak conductance and leak reversal potential, respectively.

When the membrane potential exceeds the threshold 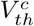, a spike is emitted and 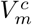 is reset to 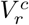, where it is held for a refractory period of 5 ms.

The noise current in the stimulus-encoding neuron 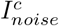 with standard deviation 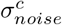 was implemented as:

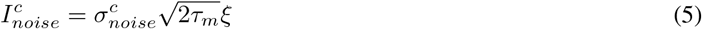

where *ξ* is Gaussian noise with zero mean and unit s.d., and *τ*_*m*_ is the membrane time constant.

A total of 3000 such neurons made synaptic connections onto the dendrites of a central auditory neuron. Parameters for the stimulus encoding model are given in Table 1. This model was written using routines from the Brian2hears package [Fontaine et al., 2011].

**Table 1:**
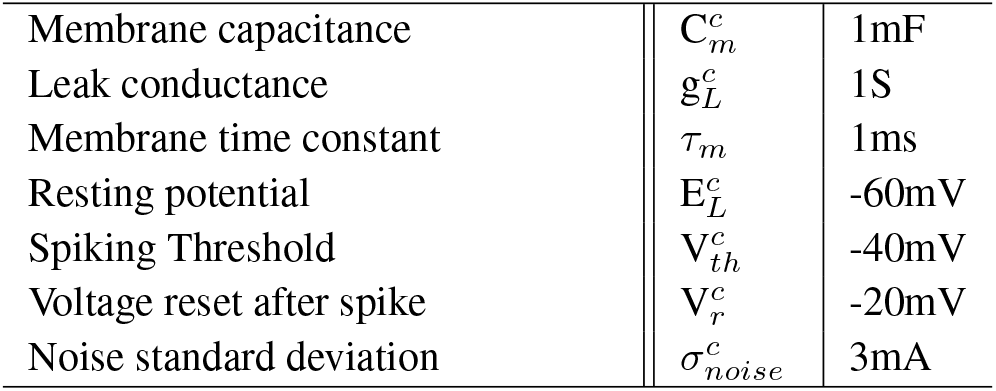
Encoding model parameters.

|  |  |  |
| --- | --- | --- |
| Membrane capacitance | $C_m^c$ | 1mF |
| Leak conductance | $g_L^c$ | 1S |
| Membrane time constant | $\tau_m$ | 1ms |
| Resting potential | $E_L^c$ | -60mV |
| Spiking Threshold | $V_{th}^c$ | -40mV |
| Voltage reset after spike | $V_r^c$ | -20mV |
| Noise standard deviation | $\sigma_{noise}^c$ | 3mA |

#### 2.5.2 Compartmental central auditory neuron

The central auditory neuron in our model consists of a soma electrically connected to 10 uniform dendritic subunits. We implemented the model using the Dendrify framework and routines [Pagkalos et al., 2023] for creating compartmental neuronal models with dendritic properties. Each compartment was modelled using an integrate-and-fire model as in Eq. 4, with inputs 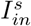 to the soma *s* or 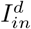 to the dendrite *d*:

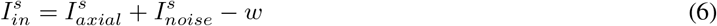

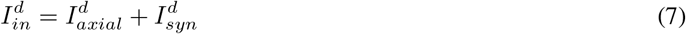

The axial current, 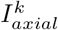, is the current that flows between connected compartments. It is a function of the voltage difference between a compartment and its neighbour, multiplied by the coupling conductance 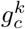. In the somatic compartment this is the sum of the currents flowing into and from all the dendritic branches denoted by *D*:

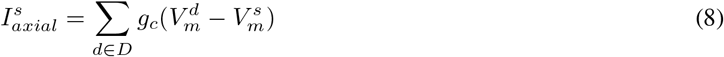

and in the dendrite, 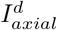 is:

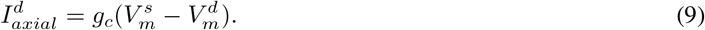

where 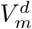 and 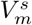 denote the dendritic and the somatic membrane potential, respectively.

An adaptive current *w* to the soma captures the transient spiking behaviour of auditory neurons recorded in the experiments. The spiking mechanism is different from that of the stimulus-encoding neuron in that when the membrane potential exceeds the threshold, 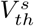, it is stepped to *V*_*spike*_ for the neuron to spike. Each time the neuron spikes, *w* is increased by a constant current, *b*, that accounts for spike-triggered adaptation. After a short delay, the membrane potential resets to 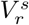. The spiking and reset events are described as follows:

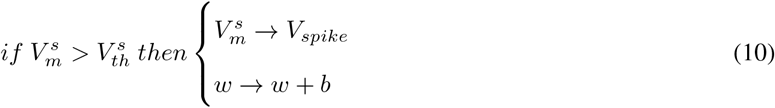

if *t* = *t*_*spike*_ + 0.2*ms* then 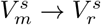.

The dynamics of *w* follow:

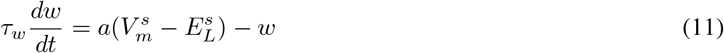

where *a* is a coupling parameter determining how sensitive the adaptation current is to the membrane potential.

The current 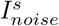 to the soma is coloured noise with standard deviation 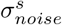, and time constant *τ*_*noise*_:

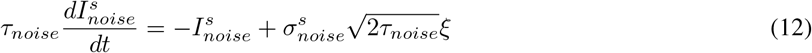

For the dendritic compartment, 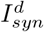 is the sum of synaptic currents entering through AMPA and NMDA receptors, where the subscript *syn* denotes the type of conductance (either AMPA or NMDA).

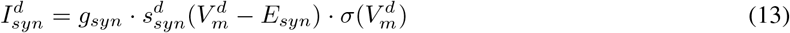

*g*_*syn*_ is the conductance for either AMPA or NMDA, *E*_*syn*_ the reversal potential of the receptor current, and *σ*(ν) describes the voltage-dependent magnesium block of the NMDA receptor:

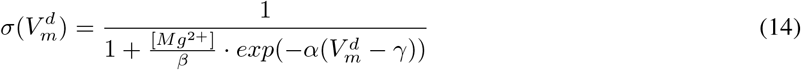

with magnesium concentration [*Mg*^2+^] = 1*mM, β* = 3.57*mM, α* = 0.062*mV* ^*−*1^ and *γ* = 0*mV* .

The time course of the synaptic conductance is governed by two equations describing the rise and decay phase with independent time constants 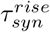 and 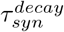:

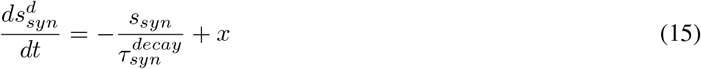

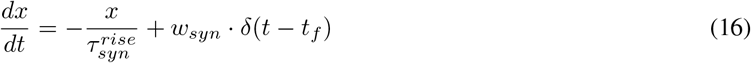

where *w*_*syn*_ is the synaptic weight and *δ*(*x*) is the Dirac delta function, and *t*_*f*_ is time of arrival of the presynaptic spike.

#### Compartment-specific properties

For a compartment with surface area *A*^*k*^, specific capacitance *c*_*m*_ and specific membrane resistivity *r*_*m*_, the membrane capacitance 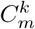 and leak conductance 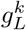 are:

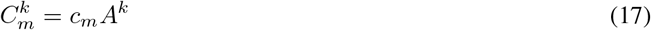

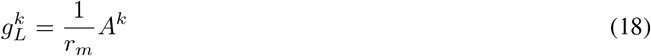

All parameters for the auditory neuron model are shown in Table 2.

**Table 2:** Central auditory neuron parameters.

|  |  |  |
| --- | --- | --- |
| Timestep | $dt$ | 0.1ms |
| Specific membrane capacitance | $c_m$ | $0.5\mu Fcm^{-2}$ |
| Specific membrane resistance | $r_m$ | $20\text{ k}\Omega cm^{-2}$ |
| Axial resistance | $r_a$ | $150\text{ }\Omega cm$ |
| Resting membrane potential | $E_L$ | -60mV |
| Spiking threshold | $V_{th}^s$ | -50mV |
| 1<sup>st</sup> voltage reset | $V_{spike}$ | 0mV |
| 2<sup>nd</sup> voltage reset | $V_r^s$ | -58mV |
| Soma length | | $25\mu m$ |
| Soma diameter | | $25\mu m$ |
| Dendrite length | | $100\mu m$ |
| Dendrite diameter | | $1\mu m$ |
| Area scale factor |  | 3 |
| Spine area factor |  | 1.5 |
| Noise time constant | $\tau_{noise}$ | 20ms |
| Noise standard deviation | $\sigma_{noise}$ | 35pA |
| Coupling conductance | $g_c$ | 0.8nS |
| Adaptation current | $b$ | 200pA |
| Adaptation coupling to membrane potential | $a$ | 0.8nS |
| Adaptation time constant | $\tau_w$ | 100ms |
| AMPA conductance | $g_{AMPA}$ | 0.8nS |
| NMDA conductance | $g_{NMDA}$ | 0.8nS |
| AMPA rise time | $\tau_{AMPA}^{rise}$ | 0.2ms |
| AMPA decay time | $\tau_{AMPA}^{decay}$ | 3ms |
| NMDA rise time | $\tau_{NMDA}^{rise}$ | 2ms |
| NMDA decay time | $\tau_{NMDA}^{decay}$ | 60ms |
| AMPA reversal potential | $E_{AMPA}$ | 0mV |
| NMDA reversal potential | $E_{NMDA}$ | 0mV |
| AMPA weight | $w_{AMPA}$ | 0.5 |
| NMDA weight | $w_{NMDA}$ | 0.05 |

#### 2.5.3 Quantifying spatial and temporal overlap on dendritic branches

We constructed spatial and temporal profiles of synaptic inputs to the dendritic branches by summing the AMPA synaptic conductance, *s*_*AMP A*_, across branches and time respectively. To quantify the degree of input overlap across branches and across time, we computed two Pearson correlation coefficients for each syllable, one between the spatial profiles and another between the temporal profiles. The median correlation value was used as a threshold to categorize inputs as dispersed and (temporally) correlated, and as clustered and uncorrelated (Figure 4B).

## 3 Results

### 3.1 MAX and AND operations in mouse auditory neurons

To probe the computations of single neurons in mouse auditory cortex, we recorded spiking responses to mouse ultrasonic vocalizations (USVs). We first identified units that were responsive to USVs and extracted syllables (*∼*100 ms in duration) coinciding with peaks in the Peristimulus Time Histogram (PSTH) (Figure 1A). Syllables extracted from the continuous USV song were presented either individually or superimposed with an appropriate time lag. The time lag was chosen to evoke the maximal response when two syllables were combined, so as not to miss the optimal temporal window for integration. For some syllable sets, the response to the combination of the two syllables was equal to or less than the response to either syllable, indicating a MAX-like response (Figure 1B). We also observed AND-like responses, where sub-threshold individual syllables would combine to evoke a response (Figure 1C), or when the superimposed syllables produced a firing rate that was greater than the arithmetic sum of the individual responses. These responses were quantified using the summation index (SmI; Eq 1). SmIs close to 0 correspond to the MAX-like operation and values equal to and greater than 1 indicate AND-like summation of inputs.

**Figure 1:**
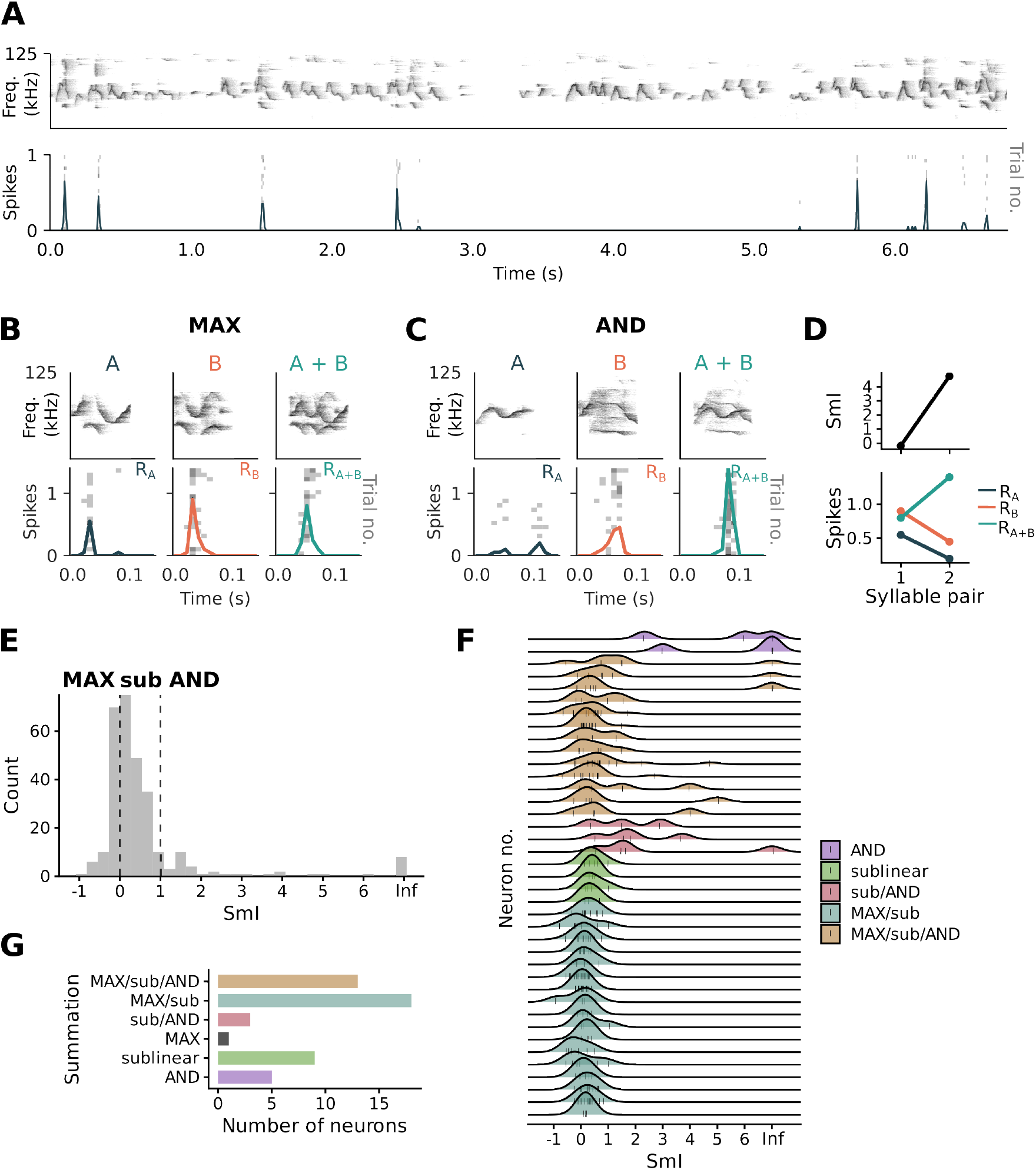
Probing feature recombination functions in single auditory neurons. (A) USV-responsive units were first identified for the experiment by presenting USV songs for several repetitions. Example PSTH (solid trace) from a single unit responding to specific USV syllables. The PSTH is overlaid over a raster plot showing spike responses over trials. (B) Pairs of syllables evoking responses in the unit were extracted and presented to the unit either separately or superimposed. Showed here are responses of the same unit in (A) to individual syllables (left and middle panels) and to their superimposition (right panel). The unit responds in a MAX-like manner to this combination of syllables. (C) Same unit showing an AND-like response to a different combination of USV syllables. (D) Firing rates to the syllables in (B) and in (C) and the resultant SmI. (E) Distribution of SmIs across all units. (F) SmIs for individual units, measured in response to different stimulus pairs. Labels correspond to the types of recombination functions exhibited by each unit. Shown here are units for which multiple SmIs were recorded. (G) A total of 49 units were recorded from auditory cortex of 21 mice. Most units exhibited more than one type of recombination function

A set of syllables activating the neuron through shared inputs would also produce a reduced response that resembles the MAX-like operation, due to input saturation. To identify truly MAX-like operations from overlapping responses, we performed an additional control, testing whether each set of syllables activated the neuron through independent inputs [Kozlov and Gentner, 2014]. Under a protocol to test cross-adaptation (Figure 2A), input independence was verified for half of the MAX responses recorded (57 of 107). For these stimuli pairs, adaptation to the repeated presentation of one syllable preserved at least 40% of the response to the other (deviant) syllable in the sequence (Figure 2C).

**Figure 2:**
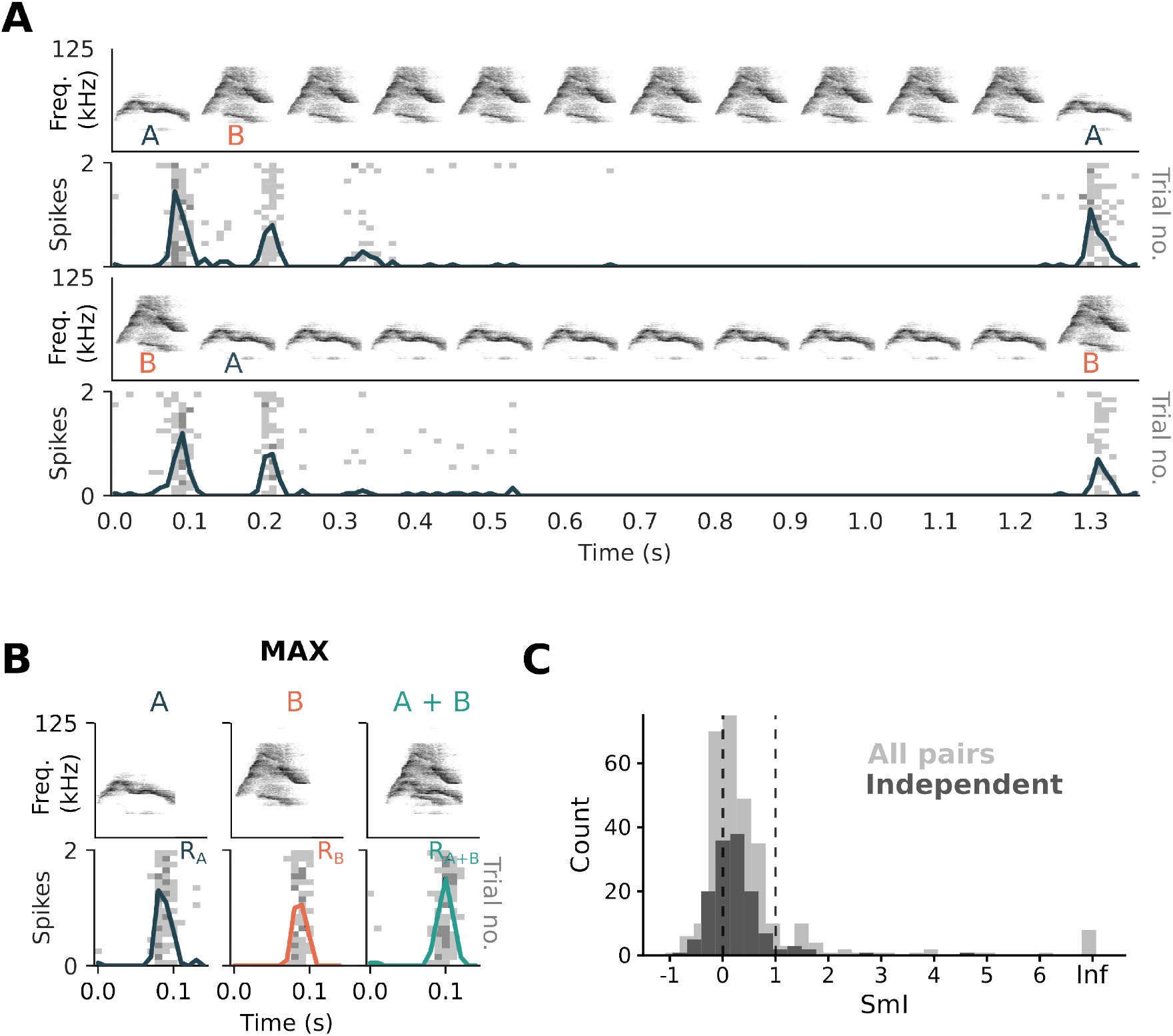
Testing input independence of pairs of syllables. (A) Cross-adaptation protocol. Example unit showing robust responses to the first syllable before and after repeated presentations of the second, suggesting that the two stimuli are activating the neuron through independent inputs. Bottom panel showing responses when the order of the two syllables was reversed. (B) Combination of the two independent syllables in (A) produced a MAX-like response by the unit. (C) Histogram of SmI values recorded from all sets of stimuli (light grey) and for pairs of independent stimuli (dark grey).

### 3.2 Single-neuron flexibility in input integration

Across all units, we observed a range of summation behaviors, with the peak of the distribution of summation indices close to 0 (Figure 1E), suggesting a predominance of the MAX-operation in this population of cells. Individual units exhibited a range of summation behaviours for different syllable pairs (Figure 1F) and were capable of implementing more than one type of recombination function (Figure 1G). To investigate changes in the neuron’s integration behaviour to simple feature transformations, we pitch-shifted (up and down by 6 semitones) syllables of each pair (Supplementary Figure 1A). Some units were broadly tuned and retained responses to the pitch-shifted syllables (Supplementary Figure 2A), while other units responded selectively to syllables pitch-shifted within a limited range (Supplementary Figure 2B). Combining pitch-shifted versions of the same base pair of syllables altered the neuronal SmI (Supplementary Figure 1D). Hence a neuron’s summation behavior, as measured by the SmI, changes in an input-dependent manner, highlighting the flexibility of single neurons in implementing different integration operations.

### 3.3 Factors influencing stimuli integration

As the integration of USV syllables spanned the sublinear to supralinear regimes, we wondered what factors determined the SmI. The arithmetic sum of responses to individual syllables (Ra + Rb) correlated negatively with SmI values (Supplementary Figure 3A; Spearman’s *ρ*_(*n*=280)_ = *−*0.42, *p <* 0.001). This suggests that integration was more MAX-like with strongly activating inputs and more AND-like with weakly activating or subthreshold inputs. Recovery from cross-adaptation (the neuron’s response to a syllable after adapting to the other syllable in the pair) correlated weakly with SmI (Supplementary Figure 3B; Spearman’s *ρ*_(*n*=280)_ = 0.18, *p* = 0.003), with more independent inputs eliciting a higher, more AND-like, SmI.

To examine if tuning properties of the neuron affected SmI, we characterized the selectivity of neuronal responses to pitch-shifting using a sparsity index (Eq 2). For each pair of syllables contributing to the SmI, two sparsity indices were calculated, each representing the degree of selectivity to pitch-shifted versions of the respective syllable (Supplementary Figure 2C). A positive correlation was observed between SmI and pitch selectivity, which was more pronounced for the smaller (less sparse) index in each pair (Supplementary Figure 2D; *ρ*_(*n*=183)_ = 0.35, *p <* 0.001) compared to the higher (more sparse) index (*ρ*_(*n*=183)_ = 0.25, *p <* 0.001). Integration tends to be more AND-like when the neuron exhibits greater pitch selectivity for a given syllable.

Besides response properties, we examined if acoustical properties of the stimuli could affect a neuron’s SmI. Specifically, we were interested in whether feature integration was biased by how acoustically distinct the stimuli were, which would have implications for object recognition computations. We characterized syllable dissimilarity using two metrics, calculating the distance between either (i) flattened spectrograms or (ii) feature vectors consisting of syllable spectral properties (frequency quartiles, mean frequency and bandwidth).

Comparing MAX-like, sublinear and AND-like SmIs, there was an overall difference in the degree of syllable dissimilarity as measured by spectrogram distance (Supplementary Figure 3C; non-parametric ANOVA with observations grouped by neuron, *F*_(2,283.6)_ = 4.16, *p* = 0.017). Post-hoc analyses with Tukey’s pairwise-comparison test showed that this difference was between MAX-like and sublinear SmIs (*t*_(250)_ = *−*2.796, *p* = 0.015), indicating that the MAX-like operation may be employed when inputs are more similar in spectrotemporal content. Although the distance between spectrogram pairs producing AND-like SmIs appeared to be greater than that of MAX-like SmIs, this difference was not statistically significant (*t*_(277)_ = *−*1.069, *p* = 0.53).

A traditional approach for comparing and characterizing vocalizations is to extract certain acoustic features from the stimulus waveform [Elie and Theunissen, 2018]. Syllables were represented using feature vectors and the distance between vectors of a pair was used as the second metric of syllable dissimilarity. As sensory coding models [Rust et al., 2005] assume that spiking is determined by recent stimuli, spectral features were extracted from a 55-ms window of the firing response (the peak of the PSTH). We found a weak positive correlation between spectral feature distance and SmI (Supplementary Figure 3D; *ρ*_(*n*=280)_ = 0.22, *p <* 0.001). This relationship was more evident for pairs of stimuli that were temporally anti-correlated (i.e., correlation between power profiles over time *<* 0, see methods); *ρ*_(*n*=91)_ = 0.4, *p <* 0.001). These results indicate that the integration of dissimilar features tends towards more AND-like summation.

To assess the relative importance of these various factors influencing the SmI, we fit a mixed-effects linear regression model with response and feature properties as regressors. The coefficients of the regressors for the sum of individual responses (*R*_*a*_ + *R*_*b*_; *t*_(114)_ = *−*7.107, *p <* 0.001), spectral feature difference (*t*_(274)_ = 4.78, *p <* 0.001), and temporal correlation (*t*_(270)_ = *−*2.43, *p* = 0.016) were found to be significant. The degree of input independence (as measured by the recovery in response following the the adapting syllable) had no significant impact on the SmI (*t*_(269)_ = 1.44, *p* = 0.15). The sum of individual responses accounted for the largest proportion of variance in the model (20%), spectral feature distance and feature temporal correlation accounted for a further 12%. These results show that while the activation strength of inputs significantly affects the SmI, the degree of dissimilarity between inputs can also influence the neuron’s integration response.

### 3.4 Investigating single-neuron flexibility in a biophysical neuron model

As our experimental results showed that neuronal response properties and feature characteristics can influence the SmI, we turned to a simple model of dendritic integration to explore the mechanisms underpinning these dependencies. To approximate sound encoding in the auditory periphery, sound clips were filtered through a bank of 3000 gammatone filters with center frequencies spanning 30 kHz - 120 kHz on the ERB (Equivalent Rectangular Bandwidth) scale. The filtered sound from each channel was fed as an input current into a noisy leaky integrate-and-fire neuron, producing a spike train representation of the stimulus for a certain frequency band. Neurons encoding the stimuli were connected to an adaptive integrate-and-fire ‘central auditory’ neuron with ten dendrites. Each dendrite received synaptic contacts from 300 encoding neurons such that similar frequencies converged on the same dendrite (Supplementary Figure 4). All syllable combinations used in the experiments, superimposed with a range of time overlaps (from 0 - 20 ms), were presented to the model. This produced a distribution of SmIs (*n* = 730) with a peak at 0 corresponding to the MAX operation (Figure 4A). Selecting the optimal time overlap which produced the highest integration response for each syllable combination (*n* = 146), as was done in the experiments, revealed more AND-like responses. This shows that the temporal alignment of stimuli affects the SmI distribution and is an important step in probing neuronal integration.

In theory, the simultaneous activation of inputs spaced close together on passive dendrites yields a sublinear response, owing to the reduction in synaptic driving force from each depolarization and mutual shunting of inputs. Spatially separated inputs integrate linearly, resulting in a more enhanced response [London and Häusser, 2005]. We wondered if these summation rules derived with simple synaptic inputs result in different feature recombination functions for similar versus dissimilar stimuli. As an example, we show that two syllables with input patterns that colocalize in time and space (branch) are integrated to produce a MAX-like response (Figure 3). Increasing the spectral distance between them (by pitch-shifting) produces an AND-like response. Inputs to the model that were spatially dispersed across branches and correlated in time had higher SmI values, compared to clustered and asynchronous inputs (Kruskal-Wallis test, *χ*^2^ = 22.1, p < 0.001; Figure 4B).

**Figure 3:**
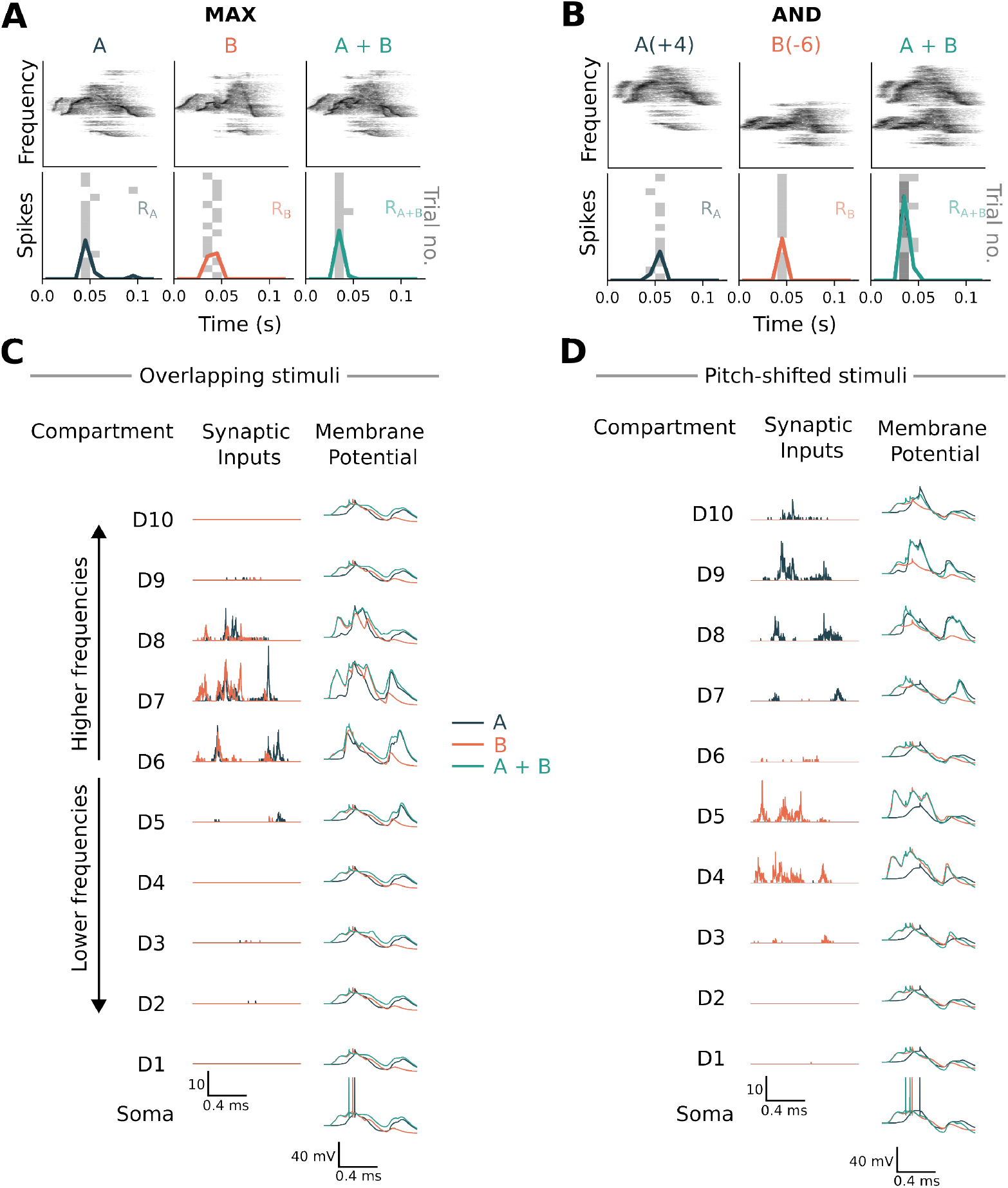
The spatiotemporal pattern of inputs on passive dendrites affects the resultant feature recombination function. (A) The MAX-like operation observed in the model for similar inputs. Top panels: Spectrograms of two syllables presented to the model. Lower panels: Spiking response of the model to two syllables, presented either in isolation or combined, over 15 trials. A small noise current injected to the soma of the neuron produces trial-to-trial variability. (B) Similar to (A) but with clip ‘A’ shifted up by 4 semitones and clip ‘B’ shifted down by 6 semitones to increase their spectral separation. This produces and AND-like response in the model. (C) Inputs that overlap in spectral content converge on the same dendrite, producing a sublinear response. Shown here are the dynamics in each compartment for the set of stimuli shown in (A). Each dendrite of the central auditory neuron receives inputs from 300 coclear filterbank neurons, with each dendrite encoding a distinct frequency range within the 30 - 120 kHz spectrum of the sound stimulus. Center column: Spatiotemporal profile of synaptic activations on each dendrite for the individual syllables. Right column: Membrane potential recorded in the dendritic and somatic compartments. The neuron emits a single spike (somatic compartment, green trace) when integrating temporally coincident and spectrally overlapping inputs. (D) Inputs separated in spectral content target different branches and are integrated linearly, leading to an enhanced response in comparison (neuron emits two spikes).

Our simulations were performed under structured connectivity, where all synapses on a given dendrite were co-tuned to a specific frequency range. Randomizing synapse locations across branches, to increase the heterogeneity of frequency tuning within individual branches, resulted in the distribution of SmIs becoming more MAX-like (Figure 4C). Furthermore, mixing inputs within each dendrite diminished the dependence of SmIs on the distance between stimuli (Figure 4C), due to a decrease in spatial segregation of distinct stimuli on dendritic branches. Thus far, all simulations were performed using a model of passive input integration. Addition of NMDA conductance shifted the distribution to more MAX-like SmIs (Figure 4D).

**Figure 4:**
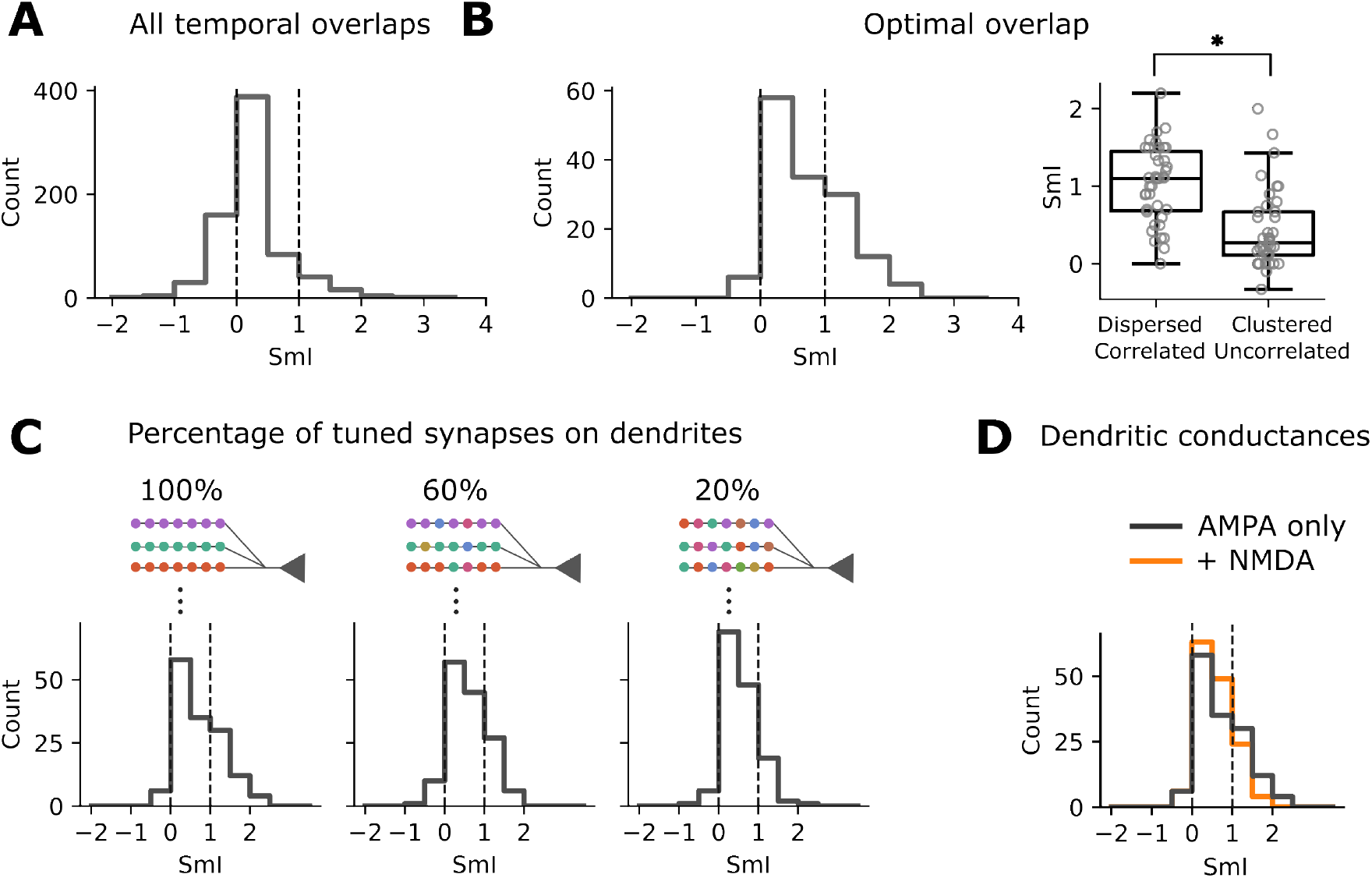
The SmI distribution changes with dendritic conductances and input mixing. (A) Distribution of SmIs for all syllable combinations, combined with different time overlaps and (B) only with the optimal time overlap, as in the experiments. (right) Boxplots of SmIs, for dispersed and temporally correlated inputs, versus clustered and uncorrelated inputs. (C) Distribution of SmIs when the degree of frequency tuning homogeneity on dendrites was varied in the passive model. We began with a structured synaptic arrangement (left), where inputs to each dendrite encode a specific frequency range of the stimulus. We then transitioned to more mixed patterns, by introducing inputs that were sampled from frequency bins outside the dendrite’s specific range. (D) Effect of adding NMDA receptors to the model on the SmI distribution.

## 4 Discussion

One can explore how neurons integrate their inputs at different levels of abstraction. At the level of biophysical mechanisms, the interplay between passive cable properties and active dendritic conductances enables nonlinear and complex transformations over synaptic inputs. On a computational level, studies have demonstrated that neurons implement the biological equivalent of logical operations integral for pattern recognition. Our work attempts to bridge these two levels of explanation. We probed the neurons’ feature recombination functions experimentally with natural and ethologically relevant stimuli to explore how it combines real-world features. We used a computational model to explore how spatiotemporal patterns of synaptic inputs on passive dendritic branches influence the resultant integration operation of the neuron.

Together with our previous study [Kozlov and Gentner, 2014], this work highlights the flexibility in single-neuron integration operations and heterogeneity across neurons. Such flexibility can arise at different stages of the mapping between stimuli and neural activity [Weber et al., 2019]. Neurons exhibit stimulus-dependent tuning and have distinct receptive fields for processing different stimulus classes [David et al., 2004, Woolley et al., 2006, Laudanski et al., 2012]. Furthermore, the nonlinearity relating receptive field-filtered inputs to firing has also been shown to be dynamic, adapting to changes in environmental statistics [Młynarski and Hermundstad, 2021, Grimes et al., 2014] and implementing normalization [Carandini and Heeger, 2011]. Here, we have focused on the input-output mapping under steady-state conditions, for instantaneous feature extraction.

When examining the impact of stimulus characteristics on SmI responses, our results showed that the integration of strongly activating features tended to be more MAX-like. In addition, we observed that increased dissimilarity between stimulus pairs biased the SmI towards more AND-like values. Further experiments are necessary to disentangle the contributions of feature activation strength and similarity. In terms of computational implications, these results align with the design of pattern recognition models. A neuron may achieve invariance to distractors by responding to a preferred stimulus which drives it strongly, thereby performing the MAX operation over its inputs. However, when features are sufficiently distinct to form a new composite, inputs may be pooled additively to achieve selectivity.

Although not studied in this work, receptive field properties are likely to influence the type of computation performed by the neuron. Currently, our biophysical model uniformly represents all stimuli across time and frequencies, neglecting tuning-specific effects on integration. In reality, the bandwidth and temporal profile of the receptive field can alter the window of integration of inputs to the neuron. Deny et al. [2017] show that the same cells perform different computations in different places of their visual field. Specifically, OFF ganglion cells switch between quasilinear and nonlinear modes of integration depending on the location of a moving bar in the receptive field. In addition, the simultaneous presentation of a bar in the receptive field center suppressed the response to a distant bar. Investigating how a neuron performs different computations for different regions of its spectrotemporal receptive field, in the context of auditory processing, presents an interesting avenue for future research.

Many studies have examined how the location, timing of inputs and dendritic conductances shape neuronal output [Cazé and Stimberg, 2021, Tran-Van-Minh et al., 2015, Makara and Magee, 2013, Branco and Häusser, 2011, Polsky et al., 2004]. For passive dendrites, spatially distributed inputs integrate linearly, resulting in an enhanced response compared to sublinear integration within dendritic branches. In contrast, clustering inputs on active dendrites engages voltage-gated conductances, producing supralinear integration [Losonczy and Magee, 2006, Polsky et al., 2004, Poirazi et al., 2003]. While these studies provided much insight into the mechanisms of neuronal integration, they were performed with well-controlled synaptic inputs. Under *in vivo* conditions, where naturalistic input patterns are complex and correlated, response predictions based on integrating simple stimuli may not hold [Ujfalussy et al., 2015, 2018] and the effect of local dendritic nonlinearities may be smaller than expected [Ujfalussy and Makara, 2020]. The relatively modest effect of increasing spectral (and hence spatial) separation distance between input stimuli on SmI suggests that there are other factors that cannot be accounted for by simple cable properties of the neuron.

In our model, synapses were highly organized into functional clusters, so that the dendrites exhibited branch-specific frequency tuning. Inputs from spectrally dissimilar stimuli were segregated over several dendrites, whereas similar features converged onto the same branches, permitting different rules of integration to operate over uncorrelated and correlated inputs. In reality, the tuning of synapses on single dendrites is likely to be more heterogeneous [Jia et al., 2010, Chen et al., 2011, Kirchner and Gjorgjieva, 2022]. More work is needed to investigate feature integration under less structured connectivity, coupled with the effect of voltage-gated dendritic conductances. Our simulations provide some indication that having an expansive nonlinearity (through the addition of NMDA receptors) in dendrites could underlie MAX-like computations, by boosting responses to individual stimuli.

Our study on single-neuron feature integration has important implications for network function and computation. The choice of activation functions in artificial neural networks can impact learning performance [Nair and Hinton, 2010, Clevert et al., 2016], yet these functions are typically static in most networks. Multiplexing computations in single neurons could unlock interesting network properties and behaviors. Adaptive input-output functions have been postulated to confer robustness to perturbations, and bring recurrent neural network dynamics to the edge of chaos, which is optimal for information propagation [Geadah et al., 2023]. Other hypothesized roles for these adaptive functions include statistical whitening transformations [Duong et al., 2023] and enabling multitask learning [Wybo et al., 2023].

In conclusion, our study has sought to investigate how neurons transform inputs to perform computations important for pattern recognition. The integration of complex spatiotemporal inputs on the dendritic tree is explored as a candidate mechanism for the single-neuron flexibility observed in the experiments. Future studies on dynamic single-neuron integration embedded in networks will provide further insights into the functional consequences of this flexibility.

## Acknowledgements

This work was funded by Biotechnology and Biological Sciences Research Council grant “How do auditory cortical neurons represent ethologically relevant natural stimuli? Characterizing stimulus feature selectivity and invariance” (BB/N008731/1).

---

**Supplementary Figure 1:**
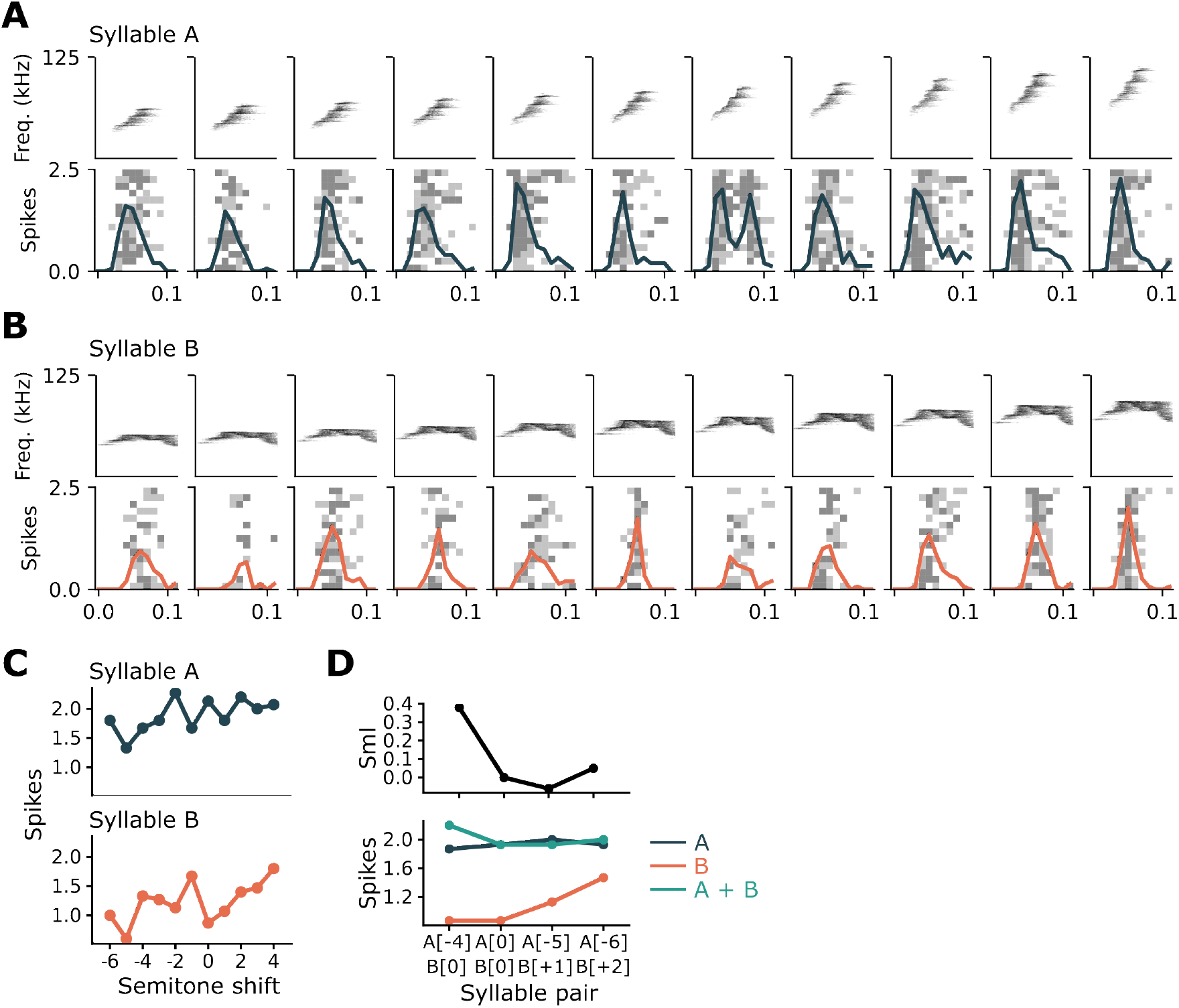
Integration of pitch-shifted inputs. (A) and (B) We tested the recombination function of a unit to different variations of the same syllable combination. Shown here are responses to pitch-shifted versions of two syllables. This unit retains good responsiveness to changes in pitch. (C) Tuning curves to pitch-shifted versions of the two syllables. (D) (Top) SmIs for pitch-shifted versions of the syllables shown in (A) and (B). (Bottom) Corresponding firing rates to individual and combined syllables; pitch-shift indicated in square brackets.

**Supplementary Figure 2:**
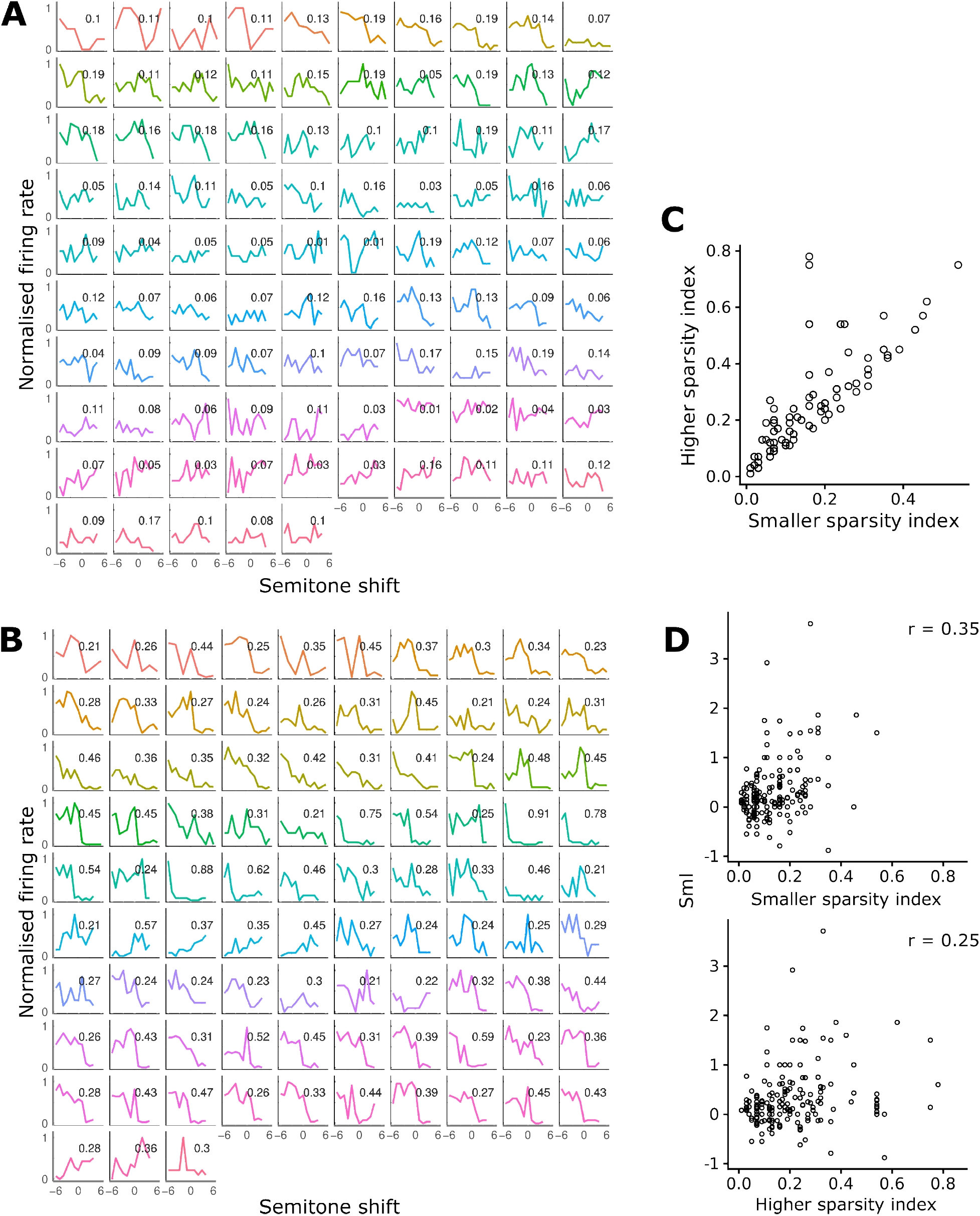
Tuning curves from pitch shifted syllables. (A) Examples of broad tuning curves (sparsity index < 0.2 where the neuron retains its response to pitch-shifting. Each panel represents tuning for a different syllable. Colors indicate different neurons; numbers indicate sparsity index. The sparsity index is a measure of the degree of selectivity of neural responses to pitch-shifted versions of a syllable. (B) Narrowly tuned pitch-selectivity profiles (sparsity index > 0.2) indicating preference for certain pitches. (C) Each point represents the sparsity indices for the two syllables in a pair, with the smaller index on the x-axis and the larger on the y-axis. (D) Correlation between sparsity index (smaller value on top) and SmI.

**Supplementary Figure 3:**
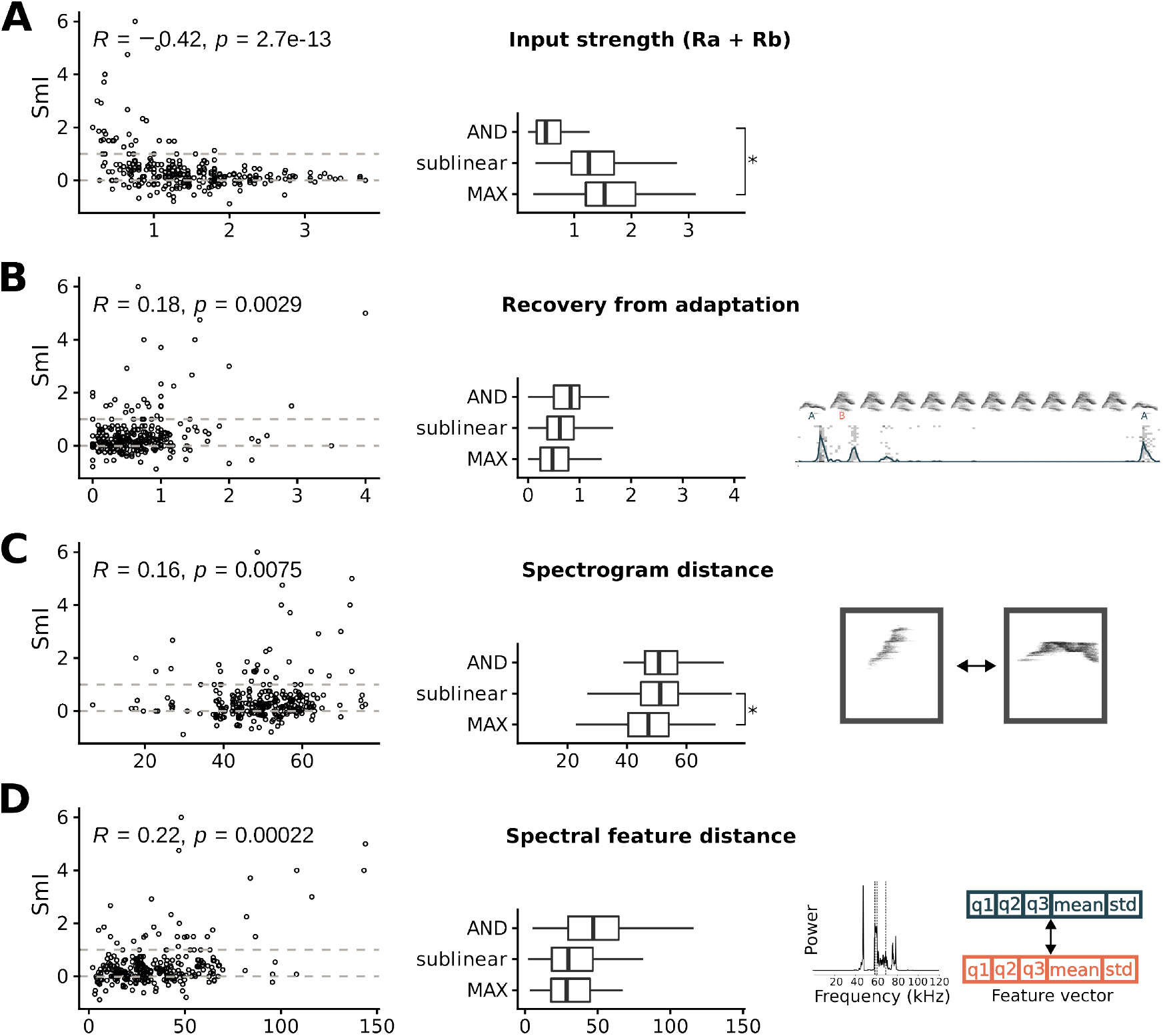
Factors influencing the summation index. (A) (Left) Scatter plot correlating input strength (the sum of responses to syllables presented individually, a proxy of how strongly the neuron is activated by input features) and SmI. (Right) Box plot comparing input strength across MAX-like, sublinear and AND-like SmIs. For statistical comparisons, SmIs from the same neuron were treated as non-independent. (B) Scatter and box plots assessing the influence of the recovery from adaptation (response to a deviant syllable after adaptation / response to the deviant syllable before adaptation). (C) Scatter and box plots showing the influence of spectrogram distance (euclidean distance between flattened spectrograms) between syllable pairs. (D) Scatter and box plots showing the influence of spectral feature difference (the distance between feature vectors representing spectral properties of the stimuli) between syllable pairs on SmI

**Supplementary Figure 4:**
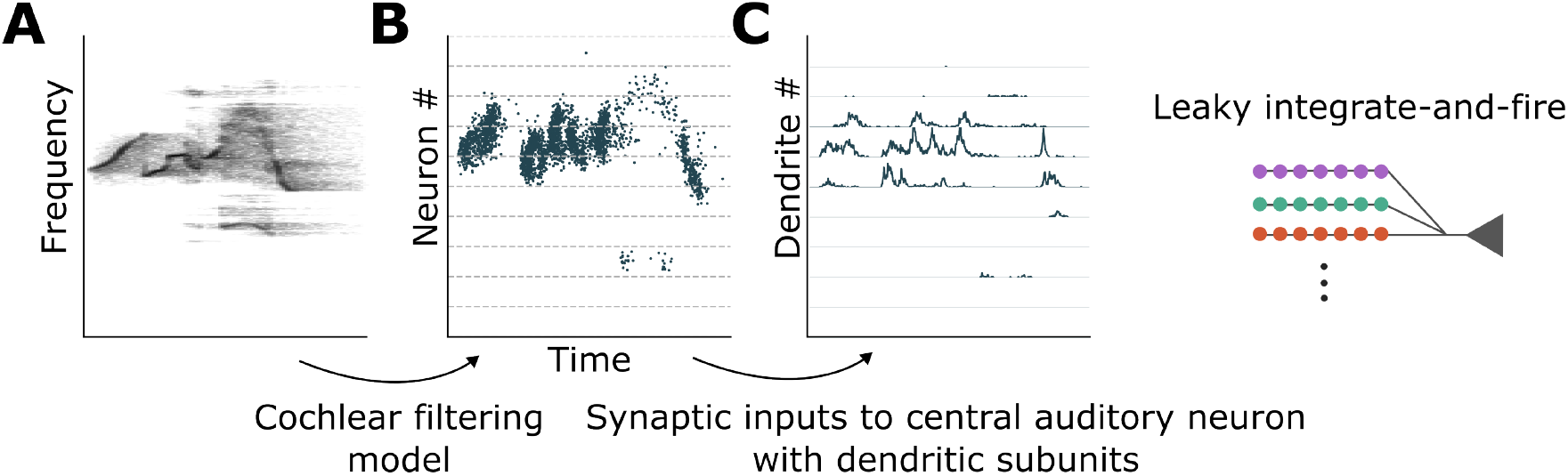
Encoding sound stimuli into spike trains. (A) Example spectrogram of a USV clip. (B) Spike train representation of the sound stimulus after processing with the auditory periphery model. Neurons with a smaller index encode low frequencies. (C) Synaptic inputs arriving at each dendrite. Each dendrite is connected to 300 neurons from (B) in the corresponding frequency band.

## References

D. Bölinger and T. Gollisch. Closed-loop measurements of iso-response stimuli reveal dynamic nonlinear stimulus integration in the retina. Neuron, 73(2):333–346, 2012. doi:10.1016/j.neuron.2011.10.039.

T. Branco and M. Häusser. Synaptic integration gradients in single cortical pyramidal cell dendrites. Neuron, 69(5): 885–892, 2011. ISSN 08966273. doi:10.1016/j.neuron.2011.02.006.

M. Carandini and D. J. Heeger. Normalization as a canonical neural computation. (November), 2011. doi:10.1038/nrn3136.

R. D. Cazé and M. Stimberg. Neurons with dendrites can perform linearly separable computations with low resolution synaptic weights. F1000Research, 9:1174, 4 2021. doi:10.12688/f1000research.26486.3.

X. Chen, U. Leischner, N. L. Rochefort, I. Nelken, and A. Konnerth. Functional mapping of single spines in cortical neurons in vivo. Nature, 475(7357):501–505, 2011. ISSN 00280836. doi:10.1038/nature10193.

D. A. Clevert, T. Unterthiner, and S. Hochreiter. Fast and accurate deep network learning by exponential linear units (ELUs). 4th International Conference on Learning Representations, ICLR 2016 - Conference Track Proceedings, pages 1–14, 2016.

S. V. David, W. E. Vinje, and J. L. Gallant. Natural stimulus statistics alter the receptive field structure of V1 neurons. Journal of Neuroscience, 24(31):6991–7006, 2004. ISSN 02706474. doi:10.1523/JNEUROSCI.1422-04.2004.

S. Deny, U. Ferrari, E. Macé, P. Yger, R. Caplette, S. Picaud, G. Tkačik, and O. Marre. Multiplexed computations in retinal ganglion cells of a single type. Nature Communications, 8(1):1–17, 2017. ISSN 20411723. doi:10.1038/s41467-017-02159-y.

L. R. Duong, D. Lipshutz, D. J. Heeger, D. B. Chklovskii, and E. P. Simoncelli. Statistical whitening of neural populations with gain-modulating interneurons. 2023. URL http://arxiv.org/abs/2301.11955.

J. E. Elie and F. E. Theunissen. Zebra finches identify individuals using vocal signatures unique to each call type. Nature Communications, 9(1), 2018. ISSN 20411723. doi:10.1038/s41467-018-06394-9. URL 10.1038/s41467-018-06394-9.

B. Fontaine, D. F. Goodman, V. Benichoux, and R. Brette. Brian hears: Online auditory processing using vectorization over channels. Frontiers in Neuroinformatics, 5(July):1–9, 2011. ISSN 16625196. doi:10.3389/fninf.2011.00009.

V. Geadah, G. Wolf, S. Horoi, G. Kerg, and G. Lajoie. Neural networks with optimized single-neuron adaptation uncover biologically plausible regularization. bioRxiv, 2023. 10.1101/2022.04.29.489963.

A. Gidon, T. A. Zolnik, P. Fidzinski, F. Bolduan, A. Papoutsi, P. Poirazi, M. Holtkamp, I. Vida, and M. E. Larkum. Dendritic action potentials and computation in human layer 2/3 cortical neurons. Science, 367(6473):83–87, 1 2020. ISSN 0036-8075. doi:10.1126/science.aax6239. URL https://www.sciencemag.org/lookup/doi/10.1126/science.aax6239.

D. F. Goodman and R. Brette. Spike-timing-based computation in sound localization. PLoS Computational Biology, 6 (11), 2010. ISSN 15537358. doi:10.1371/journal.pcbi.1000993.

W. N. Grimes, G. W. Schwartz, and F. Rieke. The synaptic and circuit mechanisms underlying a change in spatial encoding in the retina. Neuron, 82(2):460–473, 2014. ISSN 10974199. doi:10.1016/j.neuron.2014.02.037. URL 10.1016/j.neuron.2014.02.037.

H. Jia, N. L. Rochefort, X. Chen, and A. Konnerth. Dendritic organization of sensory input to cortical neurons in vivo. Nature, 464(7293):1307–1312, 2010. ISSN 00280836. doi:10.1038/nature08947.

J. H. Kirchner and J. Gjorgjieva. Emergence of synaptic organization and computation in dendrites. Neuroforum, 28(1): 21–30, 2022. ISSN 23637013. doi:10.1515/nf-2021-0031.

M. Kouh and T. Poggio. A canonical neural circuit for cortical nonlinear operations. Neural Computation, 20(6): 1427–1451, 2008. ISSN 08997667. doi:10.1162/neco.2008.02-07-466.

A. S. Kozlov and T. Q. Gentner. Central auditory neurons display flexible feature recombination functions. Journal of Neurophysiology, 111(6):1183–1189, 2014. ISSN 15221598. doi:10.1152/jn.00637.2013.

M. Lafourcade, M. S. H. van der Goes, D. Vardalaki, N. J. Brown, J. Voigts, D. H. Yun, M. E. Kim, T. Ku, and M. T. Harnett. Differential dendritic integration of long-range inputs in association cortex via subcellular changes in synaptic AMPA-to-NMDA receptor ratio. Neuron, 110(9):1532–1546, 2022. ISSN 10974199. doi:10.1016/j.neuron.2022.01.025. URL 10.1016/j.neuron.2022.01.025.

I. Lampl, D. Ferster, T. Poggio, and M. Riesenhuber. Intracellular measurements of spatial integration and the MAX operation in complex cells of the cat primary visual cortex. Journal of Neurophysiology, 92(5):2704–2713, 2004. ISSN 00223077. doi:10.1152/jn.00060.2004.

J. Laudanski, J. M. Edeline, and C. Huetz. Differences between spectro-temporal receptive fields derived from artificial and natural stimuli in the auditory cortex. PLoS ONE, 7(11), 2012. ISSN 19326203. doi:10.1371/journal.pone.0050539.

M. London and M. Häusser. Dendritic computation. Annual Review of Neuroscience, 28:503–532, 2005. ISSN 0147006X. doi:10.1146/annurev.neuro.28.061604.135703.

A. Losonczy and J. C. Magee. Integrative Properties of Radial Oblique Dendrites in Hippocampal CA1 Pyramidal Neurons. Neuron, 50(2):291–307, 2006. ISSN 08966273. doi:10.1016/j.neuron.2006.03.016.

S. Lu, G. W. Ang, M. Steadman, and A. S. Kozlov. Composite receptive fields in the mouse auditory cortex. The Journal of Physiology, 601(18):4091–4104, 2023. ISSN 0022-3751. doi:10.1113/JP285003. URL www.biorxiv.org/content/10.1101/2021.10.13.464267v1%0Ahttps://www.biorxiv.org/content/10.1101/2021.10.13.464267v1%0Ahttps://www.biorxiv.org/content/10.1101/2021.10.13.464267v1.abstracthttps://physoc.onlinelibrary.wiley.com/doi/10.1113/JP285003.

J. K. Makara and J. C. Magee. Variable dendritic integration in hippocampal CA3 pyramidal neurons. Neuron, 80(6): 1438–1450, 2013. ISSN 10974199. doi:10.1016/j.neuron.2013.10.033. URL 10.1016/j.neuron.2013.10.033.

W. F. Młynarski and A. M. Hermundstad. Efficient and adaptive sensory codes. Nature Neuroscience, 24(7):998–1009, 2021. doi:10.1038/s41593-021-00846-0.

V. Nair and G. E. Hinton. Rectified linear units improve restricted Boltzmann machines. In Proceedings of the 27th International Conference on International Conference on Machine Learning, page 807–814, 2010. doi:10.1123/jab.2016-0355.

M. Pagkalos, S. Chavlis, and P. Poirazi. Introducing the Dendrify framework for incorporating dendrites to spiking neural networks. Nature Communications, 14(1), 2023. ISSN 20411723. doi:10.1038/s41467-022-35747-8.

P. Poirazi, T. Brannon, and B. W. Mel. Pyramidal neuron as two-layer neural network. Neuron, 37(6):989–999, 2003. ISSN 08966273. doi:10.1016/S0896-6273(03)00149-1.

A. Polsky, B. W. Mel, and J. Schiller. Computational subunits in thin dendrites of pyramidal cells. Nature Neuroscience, 7(6):621–627, 2004. ISSN 10976256. doi:10.1038/nn1253.

M. Riesenhuber and T. Poggio. Hierarchical models of object recognition in cortex. Nature Neuroscience, 2(11): 1019–1025, 1999.

N. C. Rust, O. Schwartz, J. A. Movshon, and E. P. Simoncelli. Spatiotemporal elements of Macaque V1 Receptive Fields. Neuron, 46:945–956, 2005. doi:10.1016/j.neuron.2005.05.021.

T. Sato. Interactions of visual stimuli in the receptive fields of inferior temporal neurons in awake macaques. Experimental Brain Research, 77(1):23–30, 1989. ISSN 00144819. doi:10.1007/BF00250563.

J. Schiller, Y. Schiller, G. Stuart, and B. Sakmann. Calcium action potentials restricted to distal apical dendrites of rat neocortical pyramidal neurons. Journal of Physiology, 505(3):605–616, 1997. ISSN 00223751. doi:10.1111/j.1469-7793.1997.605ba.x.

J. Schiller, G. Major, H. J. Koester, and Y. Schiller. NMDA spikes in basal dendrites. Nature, 1261(1997):285–289, 2000. ISSN 1863-9135.

M. A. Steadman and C. J. Sumner. Changes in neuronal representations of consonants in the ascending auditory system and their role in speech recognition. Frontiers in Neuroscience, 12(OCT):1–16, 2018. ISSN 1662453X. doi:10.3389/fnins.2018.00671.

A. Tran-Van-Minh, R. D. Cazé, T. Abrahamsson, L. Cathala, B. S. Gutkin, and D. A. DiGregorio. Contribution of sublinear and supralinear dendritic integration to neuronal computations. Frontiers in Cellular Neuroscience, 9 (March):1–15, 2015. ISSN 16625102. doi:10.3389/fncel.2015.00067.

A. Tran-Van-Minh, T. Abrahamsson, L. Cathala, and D. A. DiGregorio. Differential dendritic integration of synaptic potentials and calcium in cerebellar interneurons. Neuron, 91(4):837–850, 2016. doi:10.1016/j.neuron.2016.07.029.

B. B. Ujfalussy and J. K. Makara. Impact of functional synapse clusters on neuronal response selectivity. Nature Communications, 11(1):1413, 2020. doi:10.1038/s41467-020-15147-6.

B. B. Ujfalussy, J. K. Makara, T. Branco, and M. Lengyel. Dendritic nonlinearities are tuned for efficient spike-based computations in cortical circuits. eLife, 4:e10056, 2015. ISSN 2050-084X. doi:10.7554/eLife.10056.

B. B. Ujfalussy, J. K. Makara, M. Lengyel, and T. Branco. Global and multiplexed dendritic computations under In vivo-like conditions. Neuron, 100(3):579–592, 2018. doi:10.1016/j.neuron.2018.08.032.

A. I. Weber, K. Krishnamurthy, and A. L. Fairhall. Coding principles in adaptation. Annual Review of Vision Science, 5: 427–449, 2019. ISSN 23744650. doi:10.1146/annurev-vision-091718-014818.

S. M. Woolley, P. R. Gill, and F. E. Theunissen. Stimulus-dependent auditory tuning results in synchronous population coding of vocalizations in the songbird midbrain. Journal of Neuroscience, 26(9):2499–2512, 2006. ISSN 02706474. doi:10.1523/JNEUROSCI.3731-05.2006.

W. A. M. Wybo, M. C. Tsai, V. A. K. Tran, B. Illing, J. Jordan, A. Morrison, and W. Senn. NMDA-driven dendritic modulation enables multitask representation learning in hierarchical sensory processing pathways. Proceedings of the National Academy of Sciences, 120(32), 8 2023. ISSN 0027-8424. doi:10.1073/pnas.2300558120. URL https://pnas.org/doi/10.1073/pnas.2300558120.

